# Assessment of thermochromic phantoms for characterizing microwave ablation devices

**DOI:** 10.1101/2024.03.23.584886

**Authors:** Ghina Zia, Amber Lintz, Clay Hardin, Anna Bottiglieri, Jan Sebek, Punit Prakash

**Affiliations:** Department of Electrical and Computer Engineering, Kansas State University, Manhattan, KS 66506, USA

**Keywords:** Tissue mimicking phantoms, thermochromic phantoms, microwave ablation

## Abstract

**Background and Purpose:** Thermochromic gel phantoms provide a controlled medium for visual assessment of thermal ablation device performance. However, there are limited studies reporting on the comparative assessment of ablation profiles assessed in thermochromic gel phantoms against those in *ex vivo* tissue. The objective of this study was to compare microwave ablation zones in a thermochromic tissue mimicking gel phantom and *ex vivo* bovine liver, and to report on measurements of the temperature dependent dielectric and thermal properties of the phantom.

**Methods:** Thermochromic polyacrylamide phantoms were fabricated following a previously reported protocol. Phantom samples were heated to temperatures in the range of 20 – 90 °C in a temperature-controlled water bath, and colorimetric analysis of images of the phantom taken after heating were used to develop a calibration between color changes and temperature to which the phantom was heated. Using a custom, 2.45 GHz water-cooled microwave ablation antenna, ablations were performed in fresh *ex vivo* liver and phantoms using 65 W applied for 5 min or 10 min (*n =* 3 samples in each medium for each power/time combination). Broadband (500 MHz – 6 GHz) temperature-dependent dielectric and thermal properties of the phantom were measured over the temperature range 22 – 100 °C.

**Results:** Colorimetric analysis showed that the sharp change in gel phantom color commences at a temperature of 57 °C. Short and long axes of the ablation zone in the phantom (as assessed by the 57 °C isotherm) for 65 W, 5 min ablations were aligned with extents of the ablation zone observed in *ex vivo* bovine liver. However, for the 65 W, 10 min setting, ablations in the phantom were on average 23.7% smaller in short axis and 7.4 % smaller in long axis than those observed in *ex vivo* liver. Measurements of the temperature dependent relative permittivity, thermal conductivity, and volumetric heat capacity of the phantom largely followed similar trends to published values for *ex vivo* liver tissue.

**Conclusion:** Thermochromic tissue mimicking phantoms provide a controlled, and reproducible medium for comparative assessment of microwave ablation devices and energy delivery settings, though ablation zone size and shapes may not accurately represent ablation sizes and shapes observed in *ex vivo* liver tissue under similar conditions.

## INTRODUCTION

Thermal ablation is a local intervention that aims to treat biological tissue by elevating tissue temperature to cytotoxic levels (> 50 °C). Ablation systems are in clinical use across a range of indications, including treatment of cardiac arrhythmias [1], [2], neuromodulation [3], solitary or oligometastatic solid tumors [4], tissue reshaping [5], and others. An ablation procedure involves localized delivery of energy to the targeted anatomy. The increase of the temperature results from the absorption of energy within the tissue. At temperatures above ∼60 °C, most tissues can be considered ablated within a few seconds [6]. The extent of the ablation zone is a function of the spatial and temporal profiles of energy distribution and absorption within tissue, subsequent passive thermal diffusion, and the cooling effects of blood perfusion [7]. Systems employing a range of energy modalities, including lasers, microwaves, radiofrequency current, and ultrasound, have been developed and are in clinical use [8].

During the design and development of ablation devices, pre-clinical assessment, and training procedures, it is desirable to have a reliable, controlled medium for evaluating device performance and spatial extents of the ablation zone. *Ex vivo* animal tissues – most frequently liver, lung, and muscle – are a widely used substrate for evaluation of ablation zone extents as a function of applied power and ablation duration, and manufacturer provided instructions for use commonly include these data [9]. The ablation zone boundary is estimated based on the extent of visible tissue discoloration, thus providing a simple method for assessing dimensions of the ablation zone; the sharpness of the ablation boundaries can be augmented through the use of viability stains when using freshly excised tissue [10], [11]. However, it can be cumbersome to obtain freshly excised tissue, and care needs to be taken to preserve tissue samples following excision to maintain sample integrity, and limit desiccation which can considerably alter tissue physical properties [12]. Additionally, variations in tissue composition (e.g. fat islets, major blood vessels) across animals may yield heterogeneity and uncertainty in tissue physical properties, which may influence ablation profiles[13], [14]. While *ex vivo* tissue does not account for the heat sink effect of blood perfusion, experiments in non-perfused tissue can serve to bound the largest extent of ablation zone anticipated under *in vivo* conditions (i.e. all else equal, if blood perfusion were present, the ablation zone would likely be smaller); *ex vivo* perfused tissue models have also been developed and reported [15]. It is noted that the initial temperature of tissue (i.e. room temperature or ∼37 °C physiologic temperature) also has a considerable impact on the extent of the observed ablation zone in benchtop experiments.

Tissue mimicking phantoms serve as an important platform for the assessment of medical devices and systems, and are widely used for quality assurance of clinical imaging and therapy systems. While experiments in *ex vivo* and *in vivo* tissue may be more representative of the intended clinical use, tissue mimicking phantoms can be constructed in a repeatable and reliable manner, with material properties tailored to match the target tissue properties. In the context of thermal therapy devices, tissue mimicking phantoms (e.g., opaque agar phantoms [16], [17], [18], [19] and transparent polyacrylamide phantoms [20]) have been widely used for evaluating the spatial profile of the ablation zone by the device under evaluation, and for comparative assessment against performance predicted by computational models [21], [22]. These require the use of temperature sensors (typically invasive, such as fiber-optic sensors) to capture transient temperature profiles during power exposure. For short duration exposures (≤ ∼30 s), the rate of heating is proportional to the power absorption in the medium, thus providing a means to evaluate the spatial power absorption pattern with appropriately positioned temperature sensors [23]. The estimated spatial power absorption profile is sensitive to temperature sensor positioning due to the steep radial gradient of power absorption and temperature [23], [24]. The development of fiber-optic thermometry based on fiber bragg grating (FBG) technology has provided a means for considerably improved spatial sampling of temperature profiles, due to the ability to record temperatures at multiple points along a fiber [25]. Non-invasive, volumetric thermometry techniques such as magnetic resonance imaging (MRI) thermometry [26], can also be used, but integrating ablation instrumentation within the MRI environment is a considerable barrier.

Recent reports have presented a range of gel phantoms that provide a facile means for estimating the extent of the ablation zone based on the color change in the phantom material at a corresponding temperature. Chen *et al* [27] utilized an agarose-albumin-based phantom to visualize the ablation outcomes of an optically tracked radiofrequency needle applicator . They used the positional data of the device in computational ablation models to generate predictive models of ablation extents. William *et al* [28] presented a liquid crystal phantom embedded within a gel matrix (impedance adjusted to match myocardial tissue) for myocardial tissue ablation with color change between 50 °C (red) and 78 °C (black). Bu Lin *et al* [20] used bovine serum albumin with polyacrylamide gel during radiofrequency ablation to visualize coagulation temperature distribution in 3D space (displayed as ivory white color). Negussie *et al* [29] reported on a polyacrylamide gel phantom incorporating a thermochromic ink to provide a facile means for spatial assessment of thermal profiles. The thermochromic ink irreversibly changes color when exposed to temperatures in the range ∼55 – 70 °C for a few seconds/minutes, thus providing a means to evaluate the extent of ablation profiles after an ablative exposure. These thermo-chromic phantoms could provide a suitable tool for validation of applicator performance during translation of medical devices from academic/early-stage innovation labs to quality manufacturing systems, quality assurance of clinical devices, and for training of clinicians [30].

These phantoms have been employed for evaluating ablation profiles of focused ultrasound [31], radiofrequency [32], and microwave ablation [33] systems. Other studies have reported on the use of such phantoms for training of clinical operators and demonstration of new ablation technologies [34]. However, the extant literature provides limited information on how microwave ablation zone profiles observed in the phantom compare against ablation zones observed in established biological tissue under similar *ex vivo* conditions. Additionally, there are limited data available on the physical properties of the phantoms, which are requisite for comparative assessment of experimental measurements against computational modeling.

The overall objective of this study was to comparatively assess ablation profiles created by a 2.45 GHz microwave ablation applicator in a previously reported thermochromic tissue mimicking gel phantom [29] and freshly excised *ex vivo* bovine liver (∼ 23 °C) tissue in a benchtop experimental setup. We also report on experimental characterizations of the frequency and temperature dependent dielectric properties, and temperature dependent thermal properties of the phantom.

## MATERIALS AND METHODS

### Thermochromic phantom preparation

Phantoms were prepared according to the protocol described by Mikhail *et al* [32]. To prepare 50 ml of phantom, combine 38.05 ml distilled water (at room temperature), 8.75 ml aqueous 40% acrylamide/bis-acrylamide (19:1 feed; a9926, Sigma Aldrich), 2.5 ml magenta ink (Kromagen WB Flexo Ink Magenta K60), and 450 mg sodium chloride in a beaker using a magnetic stirrer. A micro-pipette was used for measuring out small liquid volumes. The materials were continuously stirred until the ink had thoroughly combined. Then, 70 mg ammonium persulfate (a3678, Sigma Aldrich) was added to the mixture and stirred continuously until combined. Finally, 0.07 ml N, N, N’, N’ tetramethylethlenadiamine (TEMED, t9281, Sigma Aldrich) was added to the mixture, while continuously stirring for 30 s. The mixture was then promptly poured into storage containers and stored in a refrigerator (4 °C). All phantoms were stored in a refrigerator overnight (∼12-18 hours) prior to use in any experiments.

### Baseline characterization of colorimetric change following heating

Experiments were conducted to assess colorimetric changes to the gel phantom following exposure to temperatures in the range ∼20 – 90 °C, to validate the range of temperatures over which color changes are observed. For these experiments, a total of 50 ml of phantom was prepared and individual 3 ml samples were each placed in 4 ml glass vials (i.e., one sample consists of ∼3 ml of phantom placed in a vial), and stored overnight in a refrigerator prior to use. Prior to conducting these experiments, the vials were removed from the refrigerator and allowed to equilibrate to room temperature on the bench. The vials were then placed in a temperature-controlled water-bath (PolyScience, 2L water bath) set to temperatures in the range ∼20 – 90 °C. A needle thermocouple (MT-23, Physitemp, Clifton, NJ) was placed into the phantom during these experiments to observe the temperature of the phantom. When the phantom reached the desired temperature, and was maintained at that temperature for 1 min, the vial was removed from the water bath and allowed to rest on the bench for 1 min. Next, the vials were placed on a white background in a portable Mini Photo Studio Box (SANOTO, Whittier, CA) and a 12 MP iPhone 11 camera was used to capture digital images in consistent background lighting conditions. Images were then transferred to a computer for analysis. A custom-made tool in MATLAB was utilized to perform color correction on the images (based on a background color correction algorithm) and to analyze the intensity of the images in the red (R), green (G), and blue (B) color space. For each sample, a 3x3 region of interest in the center of the phantom was carefully selected to ensure minimal impact of lighting artifact; all data reported are the mean values of the intensity within the region of interest. Measurements were performed on a total of *n* = 35 phantom samples.

### Dielectric property characterization of the thermochromic phantom

The broadband (1 – 6 GHz) temperature-dependent dielectric properties of the tissue mimicking phantom were measured using the open-ended coaxial probe method, following a protocol similar to previously published studies [35]. Briefly, dielectric property measurements were made with the Keysight 85070E slim form probe coupled with an HP 8753D Vector Network Analyzer (VNA). The probe was calibrated with measurements on three standards: open circuit, short circuit, and room temperature deionized water. Samples of the phantom were prepared and cut into cubes measuring approximately 2 cm × 2 cm × 2 cm, and placed into a custom sample holder made from copper. The sample holder was placed on the surface of a hot plate, with custom fins extending from the surface of the hot plate to the top of the sample holder, providing a means to raise the sample temperature while attempting to limit the thermal gradient across the sample. In prior studies [35], [36], [37], we have demonstrated this method (for heating the sample) achieves heating rates of approximately 0.1 °Cs^-1^, a rate that is observed within microwave ablation zones. The 85070E probe was positioned roughly in the center of the phantom, and with the aid of a lab jack, the sample was raised to the probe such that the probe extended ∼5 mm into the phantom sample. A fiber-optic temperature sensor was affixed to the 85070E probe (∼5 mm proximal from the surface) to provide a collocated estimate of the sample temperature. The hot plate surface temperature was set to 300 °C, and dielectric properties were measured at 1 °C increments over the temperature range 22 °C to 100 °C. Measurements were performed on *n* = 3 phantom samples.

Figure 1 shows the setup for measurement of temperature dependent dielectric properties of the gel phantom.

**Figure 1.**
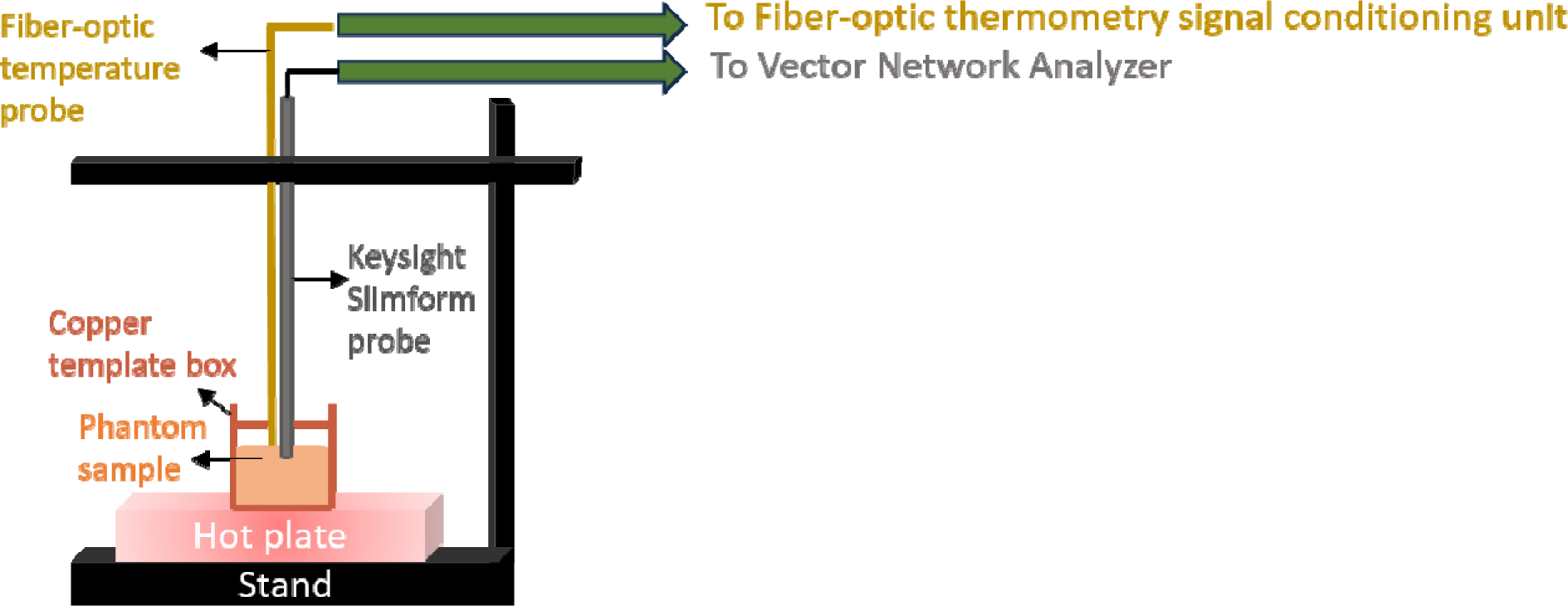
Measurement setup for temperature dependent broadband dielectric properties of the gel phantom.

### Thermal property characterization of the thermochromic phantom

Thermal properties were measured using a thermal needle system (SH-3 dual-needle, METER Group, Inc. USA) and a digital thermal analyzer (Tempos, METER Group, Inc. USA) including two parallel needles of 3 mm diameter and 30 mm length each and a 6 mm spacing between the two needles. The measurement mechanism of the system consists of a heating and a cooling phase. During the heating phase, one of the two needles is heated at a defined temperature T_0_ for about 30 s thus the heat is transferred through the sample. During the cooling phase the final temperature T_f_ is measured on the second needle. The values of thermal conductivity *k* (W ·K^-^1·m^-1^), thermal diffusivity D (m^2^· s^-1^) and volumetric heat capacity *C* (J· K^-1^·m^-3^) displayed by the system are the output parameters that minimize the difference between T_f_ and T_0_ according to the least square method.

The measurements were conducted on a total of *n* = 3 tissue mimicking phantoms exposed to a thermal controlled water bath set at a designed temperature. Each phantom was cut to fit a metallic container used to prevent the infiltration of water. The dual-needle system was placed in each sample ensuring a 15 mm margin to minimize the effect of the boundaries. The measurement procedure was initiated when the thermal equilibrium between the water bath and the phantom was reached. There were 3 phantom samples. Thermal properties were measured at 6 specific temperatures for each phantom sample within the temperature range 22-90 °C. Hence, in total the dataset consists of 18 values. Thermal property measurements involve a slight heating of the material under test by the needle-based sensor. The thermal property measurements reported here as a function of temperature refer to the baseline temperature of the phantom prior to initiating the thermal property measurement, and do not include the additional temperature change induced by the thermal property sensor.

Figure 2 shows the setup for measurement of thermal properties of the gel phantom.

**Figure 2.**
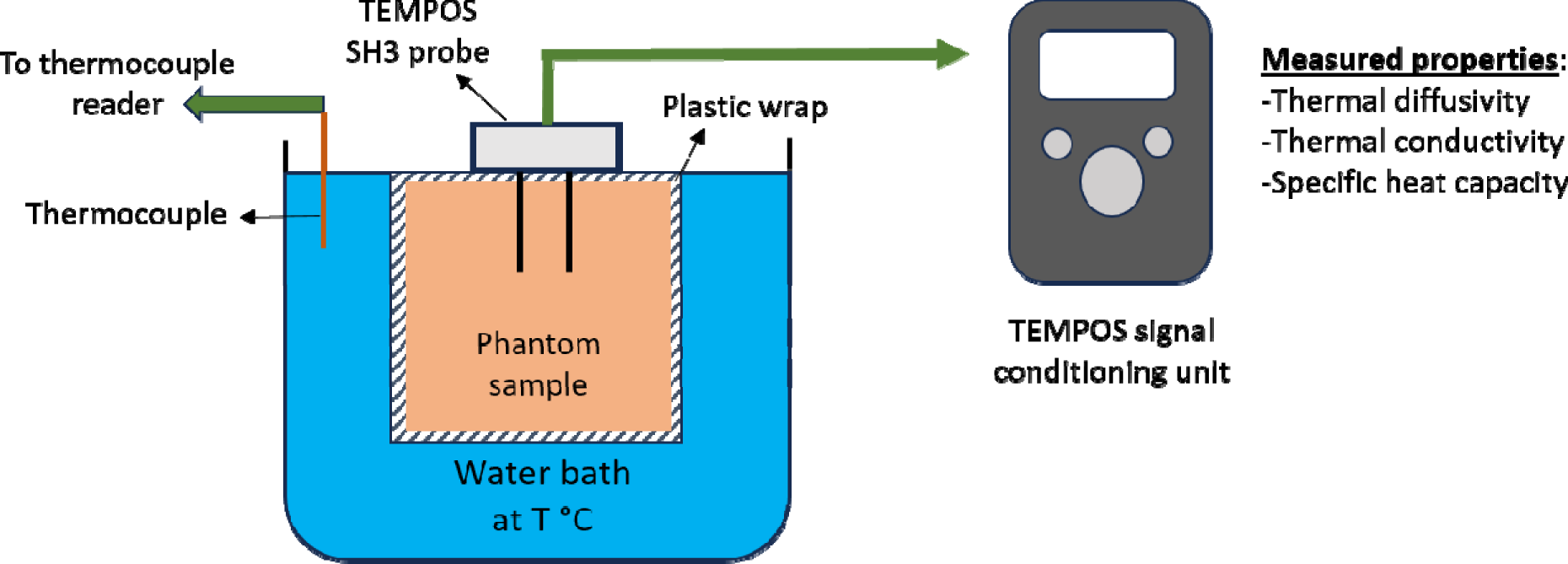
Measurement setup for thermal properties of the gel phantom

### Microwave ablation experiments in thermochromic phantoms and ex vivo bovine liver tissue

Gel phantom samples (of volume approximately 220 ml) were prepared in the lab and refrigerated. For ablation experiments, the phantoms were taken out of the refrigerator and was left on the benchtop to stabilize the temperature of the gel to room temperature (∼23 °C). The gel phantoms were placed in a custom designed acrylic template with slots positioned for arrangement of a 14 G microwave ablation applicator. Microwave ablations were performed with a custom, water-cooled monopole antenna which is similar to the one described in [38] and augmented with a 14 mm long choke (located at 20 mm from the tip of the antenna), operating at 2.45 GHz. Microwave power was supplied by a solid-state generator (Sairem GMS 200W).

Cooled water (∼7-10 °C) was circulated through the applicator using a peristaltic pump (Cole Parmer 7554–90, Vernon Hills, IL, USA) at a rate of 40 ml/min. With an applied power of 65 W, *n* = 3 experiments were performed for a heating duration of 5 min and *n* = 3 experiments were performed for a heating duration of 10 min. After the ablation experiment, the gel phantom was sliced, in a plane parallel to the applicator insertion axis, using a sharp-edged copper sheet in order to avoid disintegration of the gel phantom. Digital images of the phantoms were taken in the light-box for further analysis.

A similar procedure was followed for ablations in *ex vivo* bovine liver tissue. Briefly, freshly excised bovine liver tissue was obtained from a local slaughterhouse. Livers were placed in thin plastic bags, placed in an insulated cool box filled with ice, and transported to the lab. Within ∼4- 16 hours post excision, livers were sectioned into ∼6 cm × 6 cm × 6 cm pieces, and allowed to equilibrate to room temperature (∼23 °C) by placing the sealed Ziploc bags in a temperature-controlled water-bath. Once tissue samples had equilibrated to room temperature, they were removed from the Ziploc bags and transferred to the bench. The ablation applicator was positioned within the tissue sample and ablations were performed using applied power settings of 65 W, 5 min (*n* = 3) and 65 W, 10 min (*n* = 3). Following ablations, liver samples were sectioned in a plane parallel to the axis of insertion of the ablation applicator, and digital images of the tissue samples were taken.

## RESULTS

Figure 3 illustrates the sample sets to demonstrate color change as a function of temperature from room temperature (22 °C) to 90 °C. Each phantom vial was heated in a temperature-controlled water-bath to characterize the color of the phantom at the specific peak temperature (mentioned below the vial). The example sets of phantoms in Figure 3 indicate that from 22 °C to 55 °C peak temperature, the phantom’s light-yellow color remains consistent. The phantom starts changing its color from light-yellow to light-pink around a peak temperature of 58 °C. The phantom color gradually varies from light-pink to magenta color in range of maximum achieved sample temperatures 58 °C to 70 °C. For samples with peak temperatures above 70 °C, the color remains Magenta.

**Figure 3.**
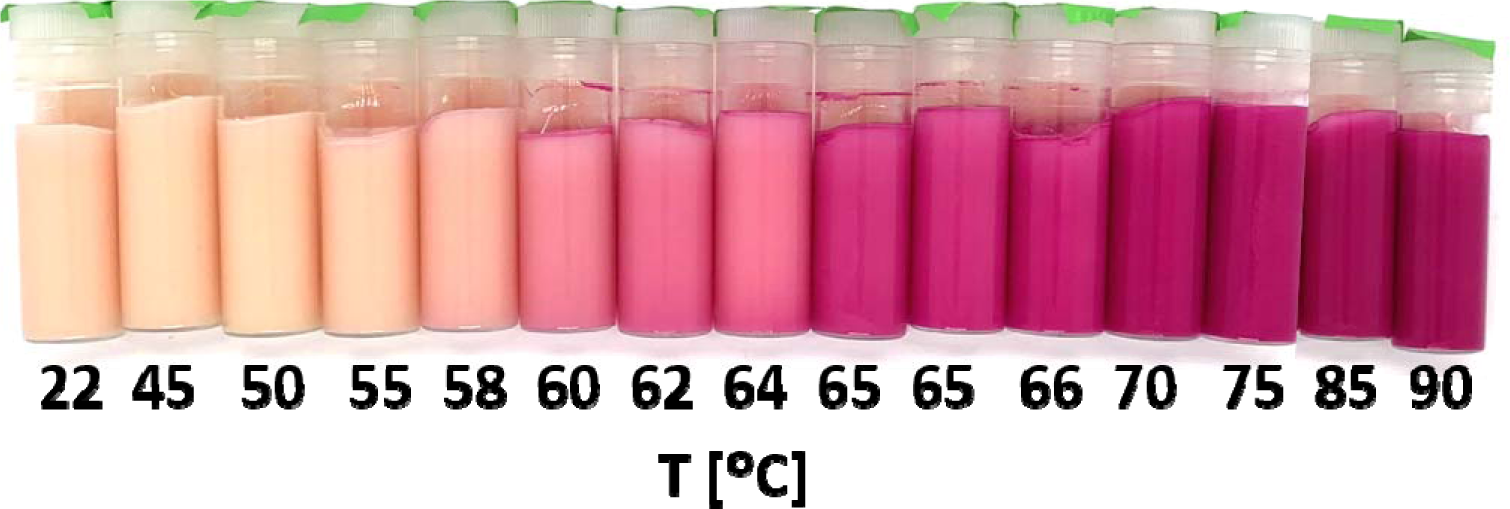
Example sets of gel phantom illustrating colorimetric changes from room temperature value (22 °C) at different peak temperature values (up to 90 °C).

Figure 4 depicts the mean RGB values obtained from the gel phantom samples after heating them to different peak temperature values. Figure 4a shows raw RGB data values from the experiments. From the RGB value analysis (Figure 4b), we determined the lowest temperature at which the color of phantom noticeably changed. This temperature and corresponding RGB values served for later ablation boundary estimation in phantom samples as well as in temperature maps predicted by the computational model. The most drastic change occurs in the green channel (“G” value) from the RGB values as compared to the red channel (or “R” values) and blue channel (or “B” values). When the phantom is exposed to peak temperatures from 57 to 72, the “G” value drops by 98% of its baseline value (at 57 peak temperature). After 72, all three color channels reach a plateau with the value of green channel being close to 0. Therefore, we used RGB values at 57 isotherm to determine ablation boundary in the phantom.

**Figure 4.**
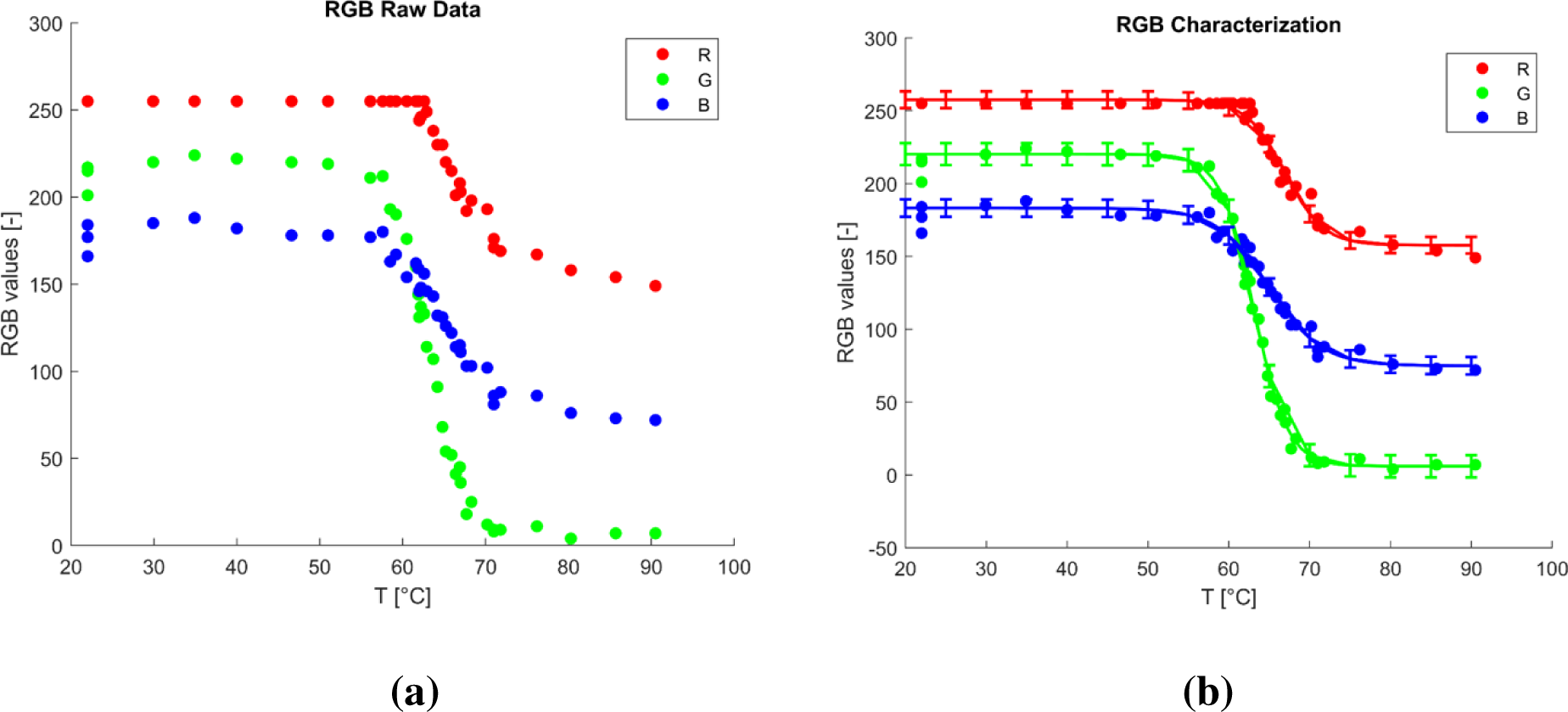
Colorimetric analysis of phantoms when exposed to temperatures in the range 20 – 90 °C in a temperature-controlled water bath. a) Raw experimental data, b) data with best-fit sigmoid curve and associated 95% confidence intervals.

For further details about the raw measurements made on each sample, please refer to the table provided in the Supplementary Materials.

Figure 5 shows example phantom and liver ablations at 65 W, 5 min, and at 65 W, 10 min. As expected, with the same power of 65 W, the 10-minute ablations in both phantom and *ex vivo* liver samples are significantly larger than the 5-minute ablations. Tables 1 and Table 2 present the dimensions of the ablations recorded in phantom and liver samples at 65 W, 5 min, and 65 W, 10 min, respectively. We analyzed images of the phantom in MATLAB to measure the width and height of the 57 peak temperature contour representing the phantom ablation dimensions. We used tissue discoloration in *ex vivo* liver samples to measure the width and height of the ablation areas.

**Figure 5.**
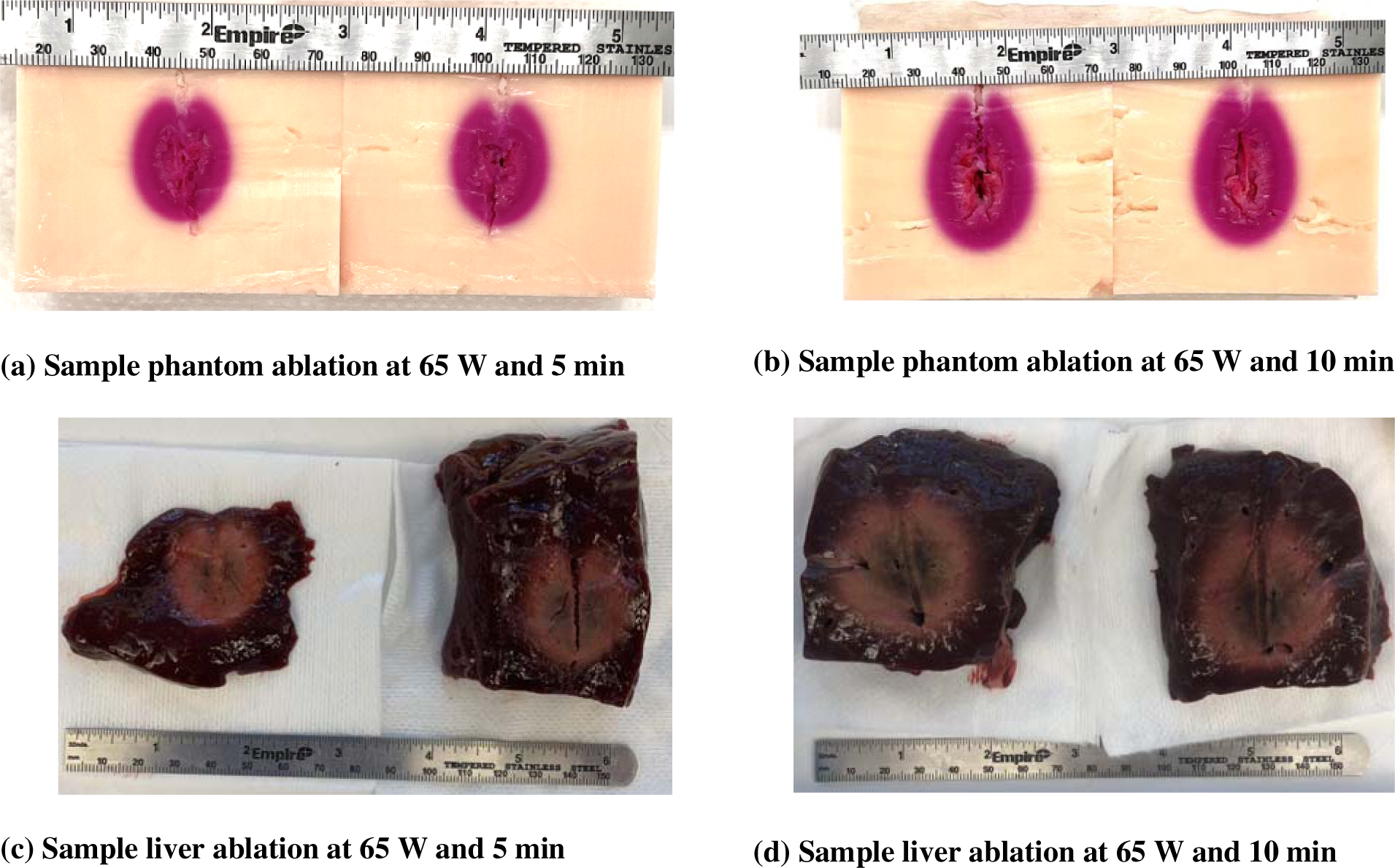
Comparison of (a) phantom and (c) *ex vivo* liver following 65 W, 5 min ablation. Comparison of (b) phantom and (d) *ex vivo* liver following 65 W, 10 min ablation.

**Table 1.** Ablation zone dimensions in phantom and liver at 65 W, 5 min.

| Sample | Ablation dimensions [cm × cm] | Average width of ablation [cm] | Average height of ablation [cm] |
| --- | --- | --- | --- |
| Phantom (Sample 1) | $2.5 \times 3.2$ | $2.4 \pm 0.1$ | $2.9 \pm 0.3$ |
| Phantom (Sample 2) | $2.3 \times 2.9$ | | |
| Phantom (Sample 3) | $2.3 \times 2.7$ | | |
| Liver (Sample 1) | $2.2 \times 3.2$ | $2.3 \pm 0.3$ | $3.0 \pm 0.1$ |
| Liver (Sample 2) | $2.1 \times 2.9$ | | |
| Liver (Sample 3) | $2.7 \times 3.0$ | | |

**Table 2.** Ablation dimensions in phantom and liver at 65 W, 10 min.

| Sample | Ablation dimensions [cm × cm] | Average width of ablation [cm] | Average height of ablation [cm] |
| --- | --- | --- | --- |
| Phantom (Sample 1) | $2.9 \times 3.9$ | $2.9 \pm 0.1$ | $3.8 \pm 0.1$ |
| Phantom (Sample 2) | $2.9 \times 3.8$ | | |
| Phantom (Sample 3) | $2.8 \times 3.7$ | | |
| Liver (Sample 1) | $4.0 \times 4.1$ | $3.8 \pm 0.3$ | $4.1 \pm 0.1$ |
| Liver (Sample 2) | $4.0 \times 4.1$ | | |
| Liver (Sample 3) | $3.5 \times 4.0$ | | |

Table 1 lists the ablation dimensions, average width and height of 65W 5 min ablation in phantom and *ex vivo* liver tissue samples. The average width of the phantom ablations is 2.4 ± 0.1 cm, which lies in the range of the average width of *ex vivo* liver tissue ablation, i.e., 2.3 ± 0.3 cm. The average height of the phantom ablations is 2.9 ± 0.3 cm, which lies in the range of the average width of *ex vivo* liver tissue ablation, i.e., 3.0 ± 0.1.

Table 2 lists the ablation dimensions, average width and height of 65 W, and 10 min ablation in phantom and *ex vivo* liver tissue samples. The average width of the phantom ablations is 2.9 ± 0.1 cm, which is around 1 cm smaller than the average width of *ex vivo* liver tissue ablation, i.e., 3.8 ± 0.3 cm. The average height of the phantom ablations is 3.8 ± 0.1 cm, which is around 0.3 cm smaller than the average width of *ex vivo* liver tissue ablation, i.e., 4.1 ± 0.1 cm.

Figure 6 compares the broadband dielectric properties of gel phantom at 35 °C over the frequency range of 500 MHz to 6 GHz for relative permittivity, ε_r_, and for effective conductivity, σ_eff_. For comparison, also shown are the broadband dielectric properties of *ex vivo* bovine liver assessed at the same temperature (taken from [35]).

**Figure 6.**
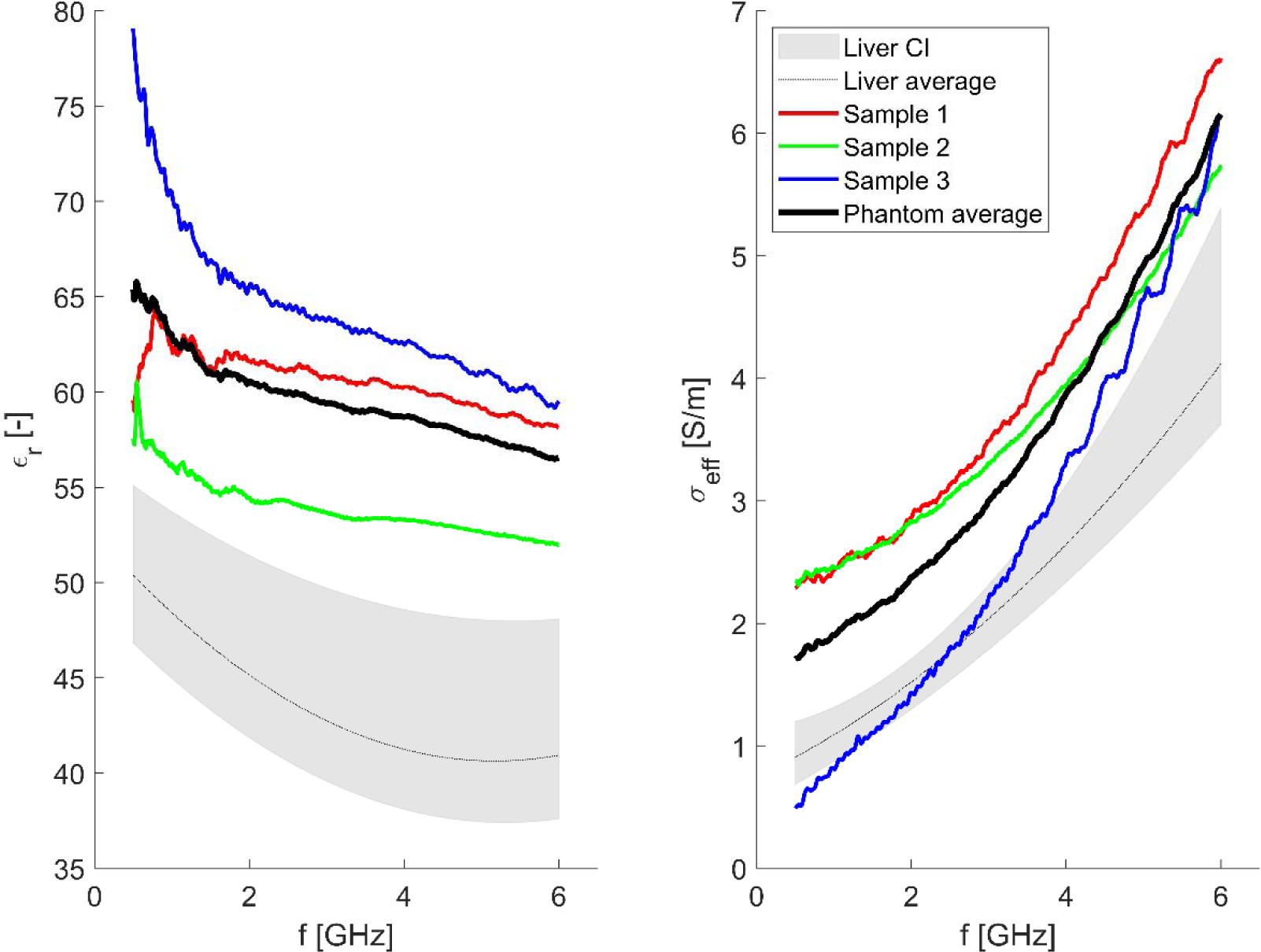
Broadband dielectric properties (relative permittivity, _r_, and effective conductivity, σ_eff_) of gel phantom at 35 °C for gel phantom (three samples with the average values) in comparison to *ex vivo* bovine liver broadband dielectric properties (average value with 95% confidence interval taken from Fallahi *et al*. 2021). **Note:** The measurement uncertainty in the broadband dielectric properties of the gel phantom is dependent on the uncertainty in the open ended coaxial probe measurements (typically within 1-5% [39], [40] https://helpfiles.keysight.com/csg/N1500A/Opt.004_Measurement_Uncertainty.htm) and the temperature sensor calibration accuracy (± 0.2°C).

Figure 7 shows the comparison of the measured relative permittivity _r_, and effective conductivity σ_eff_ of gel phantom and *ex vivo* bovine liver (taken from [35]) at Industrial, Scientific and Medical (ISM) frequencies (i.e., 915 MHz, 2.45 GHz, and 5.8 GHz) over the temperature range ∼ 22 °C to 90 °C. Several practical ablation systems have been developed for operation at ISM frequencies.

**Figure 7.**
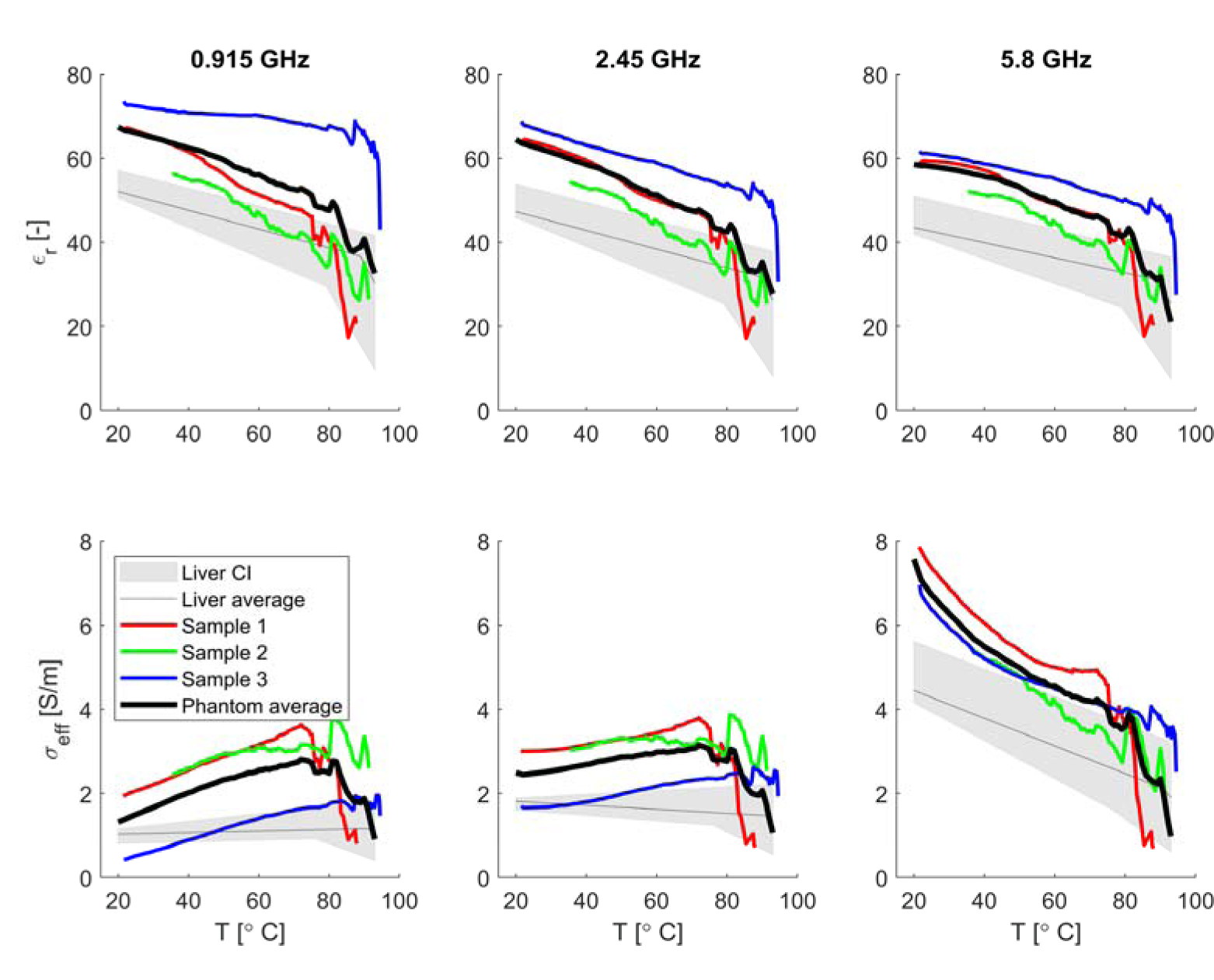
Temperature-dependent dielectric properties (relative permittivity, _r_, and effective conductivity, σ_eff_) of the gel phantom at 915 MHz, 2.45 GHz and 5.8 GHz. gel phantom (three samples with the average values) in comparison to *ex vivo* bovine liver temperature-dependent dielectric properties (average value with 95% confidence interval taken from Fallahi *et al.* 2021). Note: The measurement uncertainty in the temperature-dependent dielectric properties of the gel phantom is based on the uncertainty in the open ended coaxial probe measurements (typically within 1-5% [39], [40], https://helpfiles.keysight.com/csg/N1500A/Opt.004_Measurement_Uncertainty.htm) and the temperature sensor calibration accuracy (± 0.2°C).

Figure 8 illustrates temperature-dependent thermal properties of the gel phantom and *ex vivo* bovine liver (values taken from [41]) ; a) thermal diffusivity measured in mm^2^·s^-1^, b) Thermal conductivity measured in W.m^-1^. K^-1^, and c) volumetric heat capacity measure in MJ · m^-3^ · K^-1^, for three phantom samples and their averages over the 20-85 °C temperature range.

**Figure 8.**
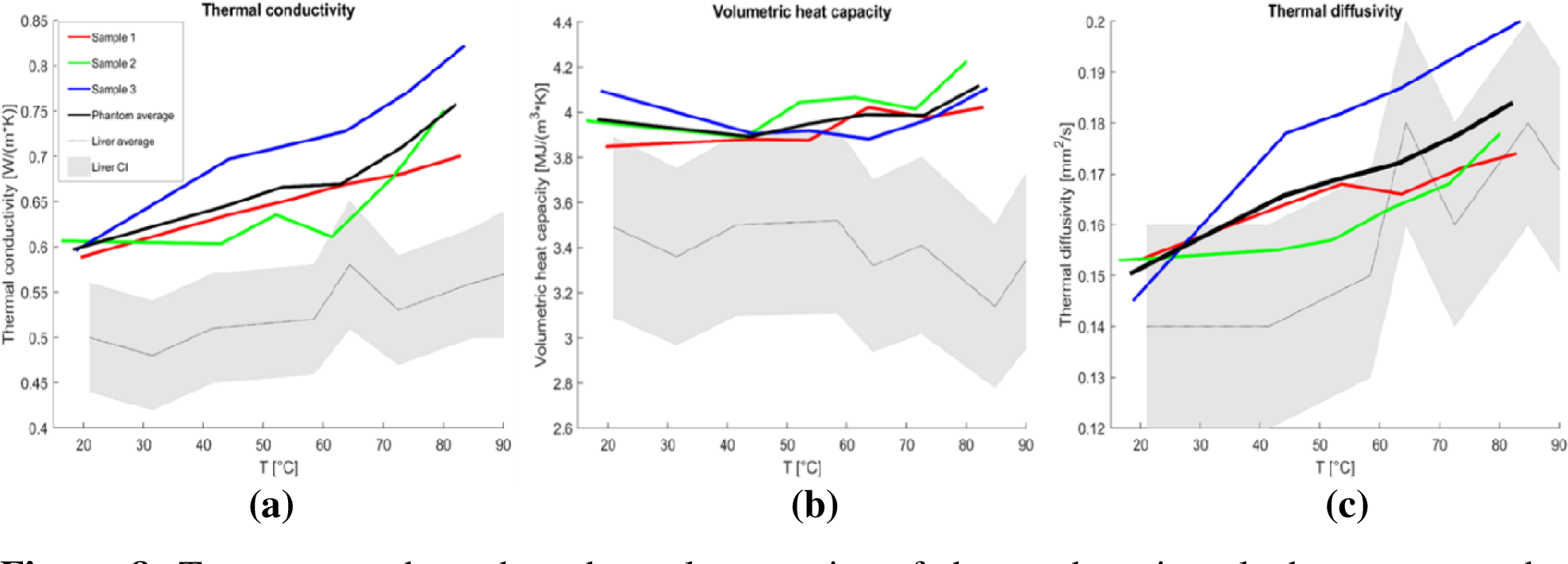
Temperature-dependent thermal properties of thermochromic gel phantom over the temperature range, 20 - 85 °C a) Thermal diffusivity, b) Thermal conductivity, and c) Volumetric heat capacity in comparison to *ex vivo* bovine liver temperature-dependent thermal properties (average with 95% confidence interval; values taken from Lopresto *et al.* 2019). **Note:** The measurement uncertainty in the temperature-dependent thermal properties of the gel phantom is dependent on the uncertainty in the Tempos SH-3 (3 cm dual needle) measurement which is ± 10% for conductivity values from 0.2–2.0 W/(m • K), for diffusivity, ±10% at conductivity > 0.2 W/ (m • K) and ±0.02 W/(m • K) for conductivities from 0.10–0.20 W/(m •) and for volumetric heat capacity, ±10% at conductivities > 0.1 W/(m • K)). (Reference: https://metergroup.com/products/tempos/tempos-tech-specs/)

## DISCUSSION

This study aimed to comparatively assess microwave ablation zones in a previously reported thermochromic phantom with ablation zones created in *ex vivo* liver when using the same ablation device and applied energy levels in both media. The colorimetric analysis was conducted to identify the working temperature range of the phantom (i.e., the range of temperatures in which the phantom significantly changes its color). The extent of the ablation zone in phantoms was determined from the 57 °C isotherm, as estimated from colorimetric analysis of phantoms heated in water baths (Figures 3 and 4). Eranki *et al* [42] conducted a similar colorimetric analysis and found the color change of the phantom was generally within the temperature range ∼ 45 °C to ∼75 °C, with the green channel displaying the most sensitivity to temperature and greatest dynamic range. In our study, we similarly found the green channel to display the greatest sensitivity to temperature change and having the greatest dynamic range. We also found no appreciable color change at temperatures below ∼57 °C (as compared to ∼ 45 °C in [42]) and above ∼72 °C (compared to ∼75 °C in [42]).

As seen in Figure 5, for 5 min ablations with 65 W applied power, similarly sized ablation zones were observed in phantoms (2.4 ± 0.1 × 2.9 ± 0.3 cm^2^) and *ex vivo* bovine liver (2.3 ± 0.3 × 3.0 ± 0.1 cm^2^). However, for 65 W, 10 min ablations, the ablation zones in phantoms (2.9 ± 0.1 × 3.8 ± 0.1 cm^2^) were on average 23.7% smaller in short axis and 7.4 % smaller in long axis than ablation zones observed in *ex vivo* liver (3.8 ± 0.3 × 4.1 ± 0.1 cm^2^). Of note, the dynamics of tissue discoloration are a function of the time-temperature history during heating, rather than temperature alone [43], whereas for the analysis of ablation zones in the phantom, the estimate of the 57 °C isotherm was used independent of duration of ablation, as in prior studies [32]. This may contribute in part to differences in ablation zone profiles observed in the phantom and liver tissue.

At higher power levels/ablation duration, the sample/tissue temperature is considerably higher than at lower applied energy settings, and the physical processes that contribute to heating and ablation zone appearance are more complex, including: vaporization and condensation carbonization, contraction, and others. These processes may contribute to the evolution of the ablation zone to varying degrees in phantoms and tissue, and thus lead to considerable differences in ablation zones observed in the two media. We hypothesized that these differences in ablation zone extents could be due in part to differences in the physical properties (dielectric properties, thermal properties, and their temperature dependencies) of the thermochromic phantom and *ex vivo* liver. We measured broadband (500 MHz – 6 GHz) dielectric properties of the phantom at temperatures ranging from ∼22 °C to ∼100 °C (Figures 6 and 7). The baseline 35 °C relative permittivity and effective conductivity values of the phantom were generally higher than that of liver tissue. At a frequency of 2.45 GHz, the relative permittivity of the phantom gradually dropped with increasing temperature, with a sharp drop at ∼ 80 °C. The effective conductivity of the phantom gradually increased at temperatures up to ∼80 °C, before a sharp drop at ∼ 80 °C. While the trend in temperature-dependent relative permittivity for the phantom at 2.45 GHz were similar to those of relative permittivity of *ex vivo* liver reported in the literature [35], [44], [45], the trend in temperature-dependent effective conductivity was somewhat different. At 2.45 GHz, the relative permittivity of liver tissue decreases slightly with increasing temperature, followed by a sharper drop at temperatures exceeding ∼80 °C; a similar trend was observed for the relative permittivity of the thermochromic phantom. At 2.45 GHz, the effective conductivity of liver tissue remains relatively stable/slightly decreases at temperatures up to ∼80 °C, and transitions to a low value at higher temperatures as tissue desiccates. This is in contrast to the slight increase in effective conductivity with temperature up to 80 °C observed for phantoms in the present study; it is noted, however, that this discrepancy may be due to the limited number of phantom samples on which measurements were conducted.

The thermal conductivity of the phantom increased with increasing temperature (Figure 8), similar to liver tissue [41], although it is noted that the baseline value of the thermal conductivity of the phantom (∼0.6 W m^-1^ K^-1^) was ∼20% higher than the value for liver at similar temperatures (∼0.5 W m^-1^ K^-1^). These larger values in thermal conductivity of the phantom relative to liver tissue (at baseline and elevated temperatures) would suggest larger zones of ablations would be observed in the phantom. However, this was not the case, possibly suggesting the role of other physical processes/factors contributing to heat transfer and color changes during microwave heating in the phantom, as compared to liver tissue. While some variations in volumetric heat capacity of the phantom were observed at varying temperatures, there was no clear trend and variation over the entire temperature range was ∼5-8 %.

The present phantom is water-based, and as such, physical properties of the phantom that change over the course of an ablation are likely linked to changes in water content during heating. For instance, Figure 5 displays regions of phantom desiccation close to the ablation applicator shaft. In Figure 7, it is seen that dielectric properties drop considerably above ∼80 °C when desiccation and vaporization may occur. Thermo-elastic modification of the phantom and other changes due to desiccation may further contribute to differences in the observed ablation zone between phantoms and *ex vivo* liver tissue.

We also note that colorimetric analyses of the phantoms were performed on samples heated uniformly in a water bath, at rates likely slower than the rate of heating during microwave ablation. Differences in the temporal evolution of color change between the calibrations obtained from colorimetric analyses compared to changes observed during heterogeneous and more rapid heating with a microwave antenna may contribute to the observed differences. It is noted that the differences in electrical properties of the phantom and liver tissue may yield differences in net power transferred from the antenna to the phantom/tissue; in order to account for this, it is recommended that future studies consider tracking forward and reflected power during ablation experiments in the phantom and liver. We also note that the initial temperature of both phantom and tissue samples in the ablation experiments was 23 °C, an accepted and widely used temperature for assessing ablation technologies on the benchtop. The energy required to raise the phantom or tissue temperature to the 57 °C threshold, would thus be more, when compared to the energy required to raise the temperature of a sample to 57 °C, when starting at physiologic temperature (i.e. ∼37 °C).

Given the increasing use of thermochromic phantoms in device evaluations and for medical simulation/training, it is important to recognize the differences in ablation zone size and shape that could be anticipated between the phantom and tissue. Further, in the *in vivo* setting, additional variability is anticipated due to the blood perfusion heat sink, and considerable variability in tissue physical properties across pathologic and normal tissue. Nevertheless, this does not diminish the utility of the phantoms, which present a controlled, stable medium for comparative assessment of ablation devices and energy delivery settings.

## CONCLUSION

The present study demonstrates that microwave ablation profiles observed in a previously reported thermochromic phantom [29] reasonably approximate ablation zones in *ex vivo* liver when for 65 W, 5 min, but not for 65 W, 10 min. Characterization of the temperature dependent thermal and dielectric properties of the thermochromic phantom material indicate differences in the baseline values of these properties compared to liver tissue, as well as temperature dependencies. These differences in physical properties may contribute to observed differences in ablation profile in the phantom and *ex vivo* liver tissue. Overall, the phantom does not appear suitable for representing liver tissue for evaluation of microwave ablation zones at high energy levels.

## ACKNOWLEDGEMENTS

This work was supported in part by NIH grants R01CA218357 and R01EB028848.

## REFERENCES

[1] D. M. Nguyen, P. Qian, T. Barry, and A. McEwan, “Cardiac radiofrequency ablation tracking using electrical impedance tomography,” Biomed Phys Eng Express, vol. 6, no. 1, p. 015015, Jan. 2020, doi: 10.1088/2057-1976/ab5ce8.

[2] P. Qian et al., “A Novel Microwave Catheter Can Perform Noncontact Circumferential Endocardial Ablation in a Model of Pulmonary Vein Isolation,” J Cardiovasc Electrophysiol, vol. 26, no. 7, pp. 799–804, Jul. 2015, doi: 10.1111/jce.12683.

[3] F. M. Gonzalez, J. Huang, and J. Fritz, “Image-Guided Radiofrequency Ablation for Joint and Back Pain: Rationales, Techniques, and Results,” Cardiovasc Intervent Radiol, vol. 46, no. 11, pp. 1538– 1550, Nov. 2023, doi: 10.1007/s00270-023-03393-2.

[4] T. P. Ryan, P. F. Turner, and and B. Hamilton, “Interstitial microwave transition from hyperthermia to ablation: Historical perspectives and current trends in thermal therapy,” Int. J. Hyperthermia, vol. 26, no. 5, pp. 415–433, 2010.

[5] E. H. Abello et al., “Temperature Profile Measurement From Radiofrequency Nasal Airway Reshaping Device,” Laryngoscope, Aug. 2023, doi: 10.1002/lary.30942.

[6] J. A. Pearce, “Models for thermal damage in tissues: processes and applications,” Crit Rev Biomed Eng, vol. 38, no. 1, pp. 1–20, 2010.

[7] D. Haemmerich, “1 - Mathematical modeling of heat transfer in biological tissues (bioheat transfer),” in *Principles and Technologies for Electromagnetic Energy Based Therapies*, P. Prakash and G. Srimathveeravalli, Eds., Academic Press, 2022, pp. 1–24. doi: 10.1016/B978-0-12-820594-5.00012-5.

[8] M. Ahmed, C. L. Brace, F. T. Lee, and S. N. Goldberg, “Principles of and advances in percutaneous ablation,” Radiology, vol. 258, no. 2, pp. 351–369, Feb. 2011, doi: 10.1148/radiol.10081634.

[9] R. Hoffmann et al., “Comparison of four microwave ablation devices: an experimental study in ex vivo bovine liver,” Radiology, vol. 268, no. 1, pp. 89–97, Jul. 2013, doi: 10.1148/radiol.13121127.

[10] J. E. Coad and J. C. Bischof, “Histologic differences between cryothermic and hyperthermic therapies,” Jun. 2003, pp. 27–36. doi: 10.1117/12.476334.

[11] B. T. Grisez, R. M. Jones, and J. E. Coad, “Evolution of pathology techniques for evaluating energy-based tissue effects,” in *Energy-based Treatment of Tissue and Assessment VII*, SPIE, Feb. 2013, pp. 150–158. doi: 10.1117/12.2006949.

[12] A. Shahzad, S. Khan, M. Jones, R. M. Dwyer, and M. O’Halloran, “Investigation of the effect of dehydration on tissue dielectric properties in ex vivo measurements,” Biomed. Phys. Eng. Express, vol. 3, no. 4, p. 045001, Jun. 2017, doi: 10.1088/2057-1976/aa74c4.

[13] J. Sebek, N. Albin, R. Bortel, B. Natarajan, and P. Prakash, “Sensitivity of microwave ablation models to tissue biophysical properties: A first step toward probabilistic modeling and treatment planning,” Medical Physics, vol. 43, no. 5, pp. 2649–2661, 2016, doi: 10.1118/1.4947482.

[14] G. Deshazer, D. Merck, M. Hagmann, D. E. Dupuy, and P. Prakash, “Physical modeling of microwave ablation zone clinical margin variance,” Medical Physics, vol. 43, no. 4, pp. 1764–1776, 2016, doi: 10.1118/1.4942980.

[15] D. Fuentes, R. Cardan, R. J. Stafford, J. Yung, G. D. Dodd, and Y. Feng, “High-fidelity computer models for prospective treatment planning of radiofrequency ablation with in vitro experimental correlation,” J Vasc Interv Radiol, vol. 21, no. 11, pp. 1725–1732, Nov. 2010, doi: 10.1016/j.jvir.2010.07.022.

[16] S. N. Goldberg, M. Ahmed, G.S. Gazelle, J.B. Kruskal, J.C. Huertas, E.F. Halpern,.. R.E. Lenkinski, “Radio-Frequency Thermal Ablation with NaCl Solution Injection: Effect of Electrical Conductivity on Tissue Heating and Coagulation—Phantom and Porcine Liver Study1,” Radiology, 219(1), 157–165, Apr. 2001, doi: 10.1148/radiology.219.1.r01ap27157.

[17] S. A. Solazzo. Z. Liu, S.M. Lobo, M. Ahmed, A.U. Hines-Peralta, R.E. Lenkinski & S.N. Goldberg, “Radiofrequency Ablation: Importance of Background Tissue Electrical Conductivity—An Agar Phantom and Computer Modeling Study1,” Radiology, 236(2), 495–502, Aug. 2005, doi: 10.1148/radiol.2362040965.

[18] Z. Liu, M. Ahmed, Y. Weinstein, M. Yi, R. L. Mahajan, and S. N. Goldberg, “Characterization of the RF ablation-induced ‘oven effect’: the importance of background tissue thermal conductivity on tissue heating,” Int J Hyperthermia, vol. 22, no. 4, pp. 327–342, Jun. 2006, doi: 10.1080/02656730600609122.

[19] D. Haemmerich and D. J. Schutt, “RF Ablation at Low Frequencies for Targeted Tumor Heating: In Vitro and Computational Modeling Results,” IEEE Transactions on Biomedical Engineering, vol. 58, no. 2, pp. 404–410, Feb. 2011, doi: 10.1109/TBME.2010.2085081.

[20] Z. Bu-Lin, H. Bing, K. Sheng-Li, Y. Huang, W. Rong, and L. Jia, “A polyacrylamide gel phantom for radiofrequency ablation,” International Journal of Hyperthermia, vol. 24, no. 7, pp. 568–576, Jan. 2008, doi: 10.1080/02656730802104732.

[21] A. Dabbagh, B. J. J. Abdullah, C. Ramasindarum, and N. H. Abu Kasim, “Tissue-mimicking gel phantoms for thermal therapy studies,” Ultrason Imaging, vol. 36, no. 4, pp. 291–316, Oct. 2014, doi: 10.1177/0161734614526372.

[22] P. Prakash, M. C. Converse, D. M. Mahvi, and J. G. Webster, “Measurement of the specific heat capacity of liver phantom,” Physiol Meas, vol. 27, no. 10, pp. N41–46, Oct. 2006, doi: 10.1088/0967-3334/27/10/N01.

[23] G. Deshazer, P. Prakash, D. Merck, and D. Haemmerich, “Experimental measurement of microwave ablation heating pattern and comparison to computer simulations,” International Journal of Hyperthermia, vol. 33, no. 1, pp. 74–82, Jan. 2017, doi: 10.1080/02656736.2016.1206630.

[24] F. Hojjatollah and P. Punit, “Antenna Designs for Microwave Tissue Ablation,” Crit Rev Biomed Eng, vol. 46, no. 6, pp. 495–521, 2018, doi: 10.1615/CritRevBiomedEng.2018028554.

[25] T. Boers et al., “An anthropomorphic thyroid phantom for ultrasound-guided radiofrequency ablation of nodules,” Med Phys, Dec. 2023, doi: 10.1002/mp.16906.

[26] P. Faridi, P. Keselman, H. Fallahi, and P. Prakash, “Experimental assessment of microwave ablation computational modeling with MR thermometry,” Medical Physics, vol. 47, no. 9, pp. 3777–3788, 2020, doi: 10.1002/mp.14318.

[27] C.-C. R. Chen, M. I. Miga, and R. L. Galloway, “Characterization of tracked radiofrequency ablation in phantom,” Medical Physics, vol. 34, no. 10, pp. 4030–4040, 2007, doi: 10.1118/1.2761978.

[28] W. W. b. Chik et al., “High Spatial Resolution Thermal Mapping of Radiofrequency Ablation Lesions Using a Novel Thermochromic Liquid Crystal Myocardial Phantom,” Journal of Cardiovascular Electrophysiology, vol. 24, no. 11, pp. 1278–1286, 2013, doi: 10.1111/jce.12209.

[29] A. H. Negussie et al., “Thermochromic tissue-mimicking phantom for optimisation of thermal tumour ablation,” International Journal of Hyperthermia, vol. 32, no. 3, pp. 239–243, Apr. 2016, doi: 10.3109/02656736.2016.1145745.

[30] A. H. Negussie, R. Morhard, J. Rivera, J. F. Delgado, S. Xu, and B. J. Wood, “Thermochromic phantoms and paint to characterize and model image-guided thermal ablation and ablation devices: a review,” Functional Composite Mater, vol. 5, no. 1, p. 1, Jan. 2024, doi: 10.1186/s42252-023-00050-2.

[31] W. Zhang et al., “Anatomic thermochromic tissue-mimicking phantom of the lumbar spine for pre- clinical evaluation of MR-guided focused ultrasound (MRgFUS) ablation of the facet joint,” Int J Hyperthermia, vol. 38, no. 1, pp. 130–135, 2021, doi: 10.1080/02656736.2021.1880650.

[32] A. S. Mikhail, A. H. Negussie, C. Graham, M. Mathew, B. J. Wood, and A. Partanen, “Evaluation of a tissue-mimicking thermochromic phantom for radiofrequency ablation,” Med Phys, vol. 43, no. 7, pp. 4304–4311, Jul. 2016, doi: 10.1118/1.4953394.

[33] N. A. Varble et al., “Morphometric characterization and temporal temperature measurements during hepatic microwave ablation in swine,” PLoS One, vol. 18, no. 8, p. e0289674, 2023, doi: 10.1371/journal.pone.0289674.

[34] X. Zhong, P. Zhou, Y. Zhao, W. Liu, and X. Zhang, “A novel tissue-mimicking phantom for US/CT/MR-guided tumor puncture and thermal ablation,” International Journal of Hyperthermia, vol. 39, no. 1, pp. 557–563, Dec. 2022, doi: 10.1080/02656736.2022.2056249.

[35] H. Fallahi, J. Sebek, and P. Prakash, “Broadband Dielectric Properties of Ex Vivo Bovine Liver Tissue Characterized at Ablative Temperatures,” IEEE Trans Biomed Eng, vol. 68, no. 1, pp. 90–98, Jan. 2021, doi: 10.1109/TBME.2020.2996825.

[36] J. Sebek and P. Prakash, “Broadband dielectric properties of porcine lung as a function of temperature,” in 2019 13th European Conference on Antennas and Propagation (EuCAP), Mar. 2019, pp. 1–5.

[37] G. Zia, J. Šebek, and P. Prakash, “Temperature-dependent dielectric properties of human uterine fibroids over microwave frequencies,” Biomed. Phys. Eng. Express, 2021, doi: 10.1088/2057-1976/ac27c2.

[38] J. Sebek, P. Taeprasartsit, H. Wibowo, W. L. Beard, R. Bortel, and P. Prakash, “Microwave ablation of lung tumors: A probabilistic approach for simulation-based treatment planning,” Med Phys, vol. 48, no. 7, pp. 3991–4003, Jul. 2021, doi: 10.1002/mp.14923.

[39] A. La Gioia et al., “Open-Ended Coaxial Probe Technique for Dielectric Measurement of Biological Tissues: Challenges and Common Practices,” Diagnostics (Basel), vol. 8, no. 2, Jun. 2018, doi: 10.3390/diagnostics8020040.

[40] C. L. Brace and S. Etoz, “An Analysis of Open-Ended Coaxial Probe Sensitivity to Heterogeneous Media,” Sensors (Basel), vol. 20, no. 18, Sep. 2020, doi: 10.3390/s20185372.

[41] V. Lopresto, A. Argentieri, R. Pinto, and M. Cavagnaro, “Temperature dependence of thermal properties of *ex vivo* liver tissue up to ablative temperatures,” Physics in Medicine & Biology, vol. 64, no. 10, p. 105016, May 2019, doi: 10.1088/1361-6560/ab1663.

[42] A. Eranki, A. S. Mikhail, A. H. Negussie, P. S. Katti, B. J. Wood, and A. Partanen, “Tissue-mimicking thermochromic phantom for characterization of HIFU devices and applications,” International Journal of Hyperthermia, vol. 36, no. 1, pp. 517–528, Jan. 2019, doi: 10.1080/02656736.2019.1605458.

[43] V. K. Nagarajan, J. M. Ward, and B. Yu, “Association of Liver Tissue Optical Properties and Thermal Damage,” Lasers Surg Med, vol. 52, no. 8, pp. 779–787, Oct. 2020, doi: 10.1002/lsm.23209.

[44] V. Lopresto, R. Pinto, G. A. Lovisolo, and M. Cavagnaro, “Changes in the dielectric properties ofex vivobovine liver during microwave thermal ablation at 2.45 GHz,” Phys. Med. Biol., vol. 57, no. 8, pp. 2309–2327, Mar. 2012, doi: 10.1088/0031-9155/57/8/2309.

[45] Z. Ji and C. L. Brace, “Expanded modeling of temperature-dependent dielectric properties for microwave thermal ablation,” Phys Med Biol, vol. 56, no. 16, pp. 5249–5264, Aug. 2011, doi: 10.1088/0031-9155/56/16/011.

