## Supplemental Table 1 for "Assessment of thermochromic phantoms for characterizing microwave ablation devices"

### Supplementary material

#### Raw RGB data values taken from gel phantom experiments with total 36 phantom samples.

| Temperature ( ºC) | R value | G value | B value |
|---|---|---|---|
| 22 | 255 | 215 | 177 |
| 22 | 255 | 201 | 166 |
| 22 | 255 | 217 | 184 |
| 29.9 | 255 | 220 | 185 |
| 34.9 | 255 | 224 | 188 |
| 40 | 255 | 222 | 182 |
| 46.6 | 255 | 220 | 178 |
| 51 | 255 | 219 | 178 |
| 56.1 | 255 | 211 | 177 |
| 57.6 | 255 | 212 | 180 |
| 58.5 | 255 | 193 | 163 |
| 59.2 | 255 | 190 | 167 |
| 60.5 | 255 | 176 | 154 |
| 61.6 | 255 | 160 | 162 |
| 61.9 | 255 | 144 | 159 |
| 62 | 244 | 131 | 146 |
| 62.2 | 246 | 137 | 148 |
| 62.6 | 255 | 133 | 156 |
| 62.9 | 249 | 114 | 146 |
| 63.7 | 238 | 107 | 143 |
| 64.2 | 230 | 91 | 132 |
| 64.8 | 230 | 68 | 131 |
| 65.2 | 220 | 54 | 126 |
| 65.9 | 215 | 52 | 122 |
| 66.4 | 201 | 41 | 114 |
| 66.9 | 208 | 45 | 115 |
| 67 | 203 | 36 | 111 |
| 67.7 | 192 | 18 | 103 |
| 68.3 | 198 | 25 | 103 |
| 70.2 | 193 | 12 | 102 |
| 71 | 176 | 9 | 81 |
| 71 | 171 | 8 | 86 |
| 71.8 | 169 | 9 | 88 |
| 76.2 | 167 | 11 | 86 |
| 80.3 | 158 | 4 | 76 |
| 85.7 | 154 | 7 | 73 |
| 90.5 | 149 | 7 | 72 |
